# Single-cell epigenomics maps the continuous regulatory landscape of human hematopoietic differentiation

**DOI:** 10.1101/109843

**Authors:** Jason D Buenrostro, M Ryan Corces, Beijing Wu, Alicia N Schep, Caleb A Lareau, Ravindra Majeti, Howard Y. Chang, William J. Greenleaf

## Abstract

Normal human hematopoiesis involves cellular differentiation of multipotent cells into progressively more lineage-restricted states. While epigenomic landscapes of this process have been explored in immunophenotypically-defined populations, the single-cell regulatory variation that defines hematopoietic differentiation has been hidden by ensemble averaging. We generated single-cell chromatin accessibility landscapes across 8 populations of immunophenotypically-defined human hematopoietic cell types. Using bulk chromatin accessibility profiles to scaffold our single-cell data analysis, we constructed an epigenomic landscape of human hematopoiesis and characterized epigenomic heterogeneity within phenotypically sorted populations to find epigenomic lineage-bias toward different developmental branches in multipotent stem cell states. We identify and isolate sub-populations within classically-defined granulocyte-macrophage progenitors (GMPs) and use ATAC-seq and RNA-seq to confirm that GMPs are epigenomically and transcriptomically heterogeneous. Furthermore, we identified transcription factors and *cis*-regulatory elements linked to changes in chromatin accessibility within cellular populations and across a continuous myeloid developmental trajectory, and observe relatively simple TF motif dynamics give rise to a broad diversity of accessibility dynamics at cis-regulatory elements. Overall, this work provides a template for exploration of complex regulatory dynamics in primary human tissues at the ultimate level of granular specificity – the single cell.

**One Sentence Summary:** Single cell chromatin accessibility reveals a high-resolution, continuous landscape of regulatory variation in human hematopoiesis.

## Main

### Introduction

In 1957, Conrad Waddington developed a fertile analogy for developmental cell biology by conceptualizing cellular differentiation as a ball rolling down a bifurcating three-dimensional surface^1,2^. This epigenomic landscape defines a descriptive pathway that a cell might follow, choosing different developmental fates as it reaches saddle points that separate different, increasingly restricted, cellular states. The shape of this epigenomic landscape is largely defined by transcription factors, which recruit chromatin effectors to reconfigure chromatin^3,4^ and promote new cellular phenotypes^5,6^. These master TF regulators represent "guy-wires” that pull on this landscape, creating troughs sufficient to attract cells into specific developmental trajectories. These concepts – the first a descriptive notion of development (**Extended Data Fig. 1a**), and the second a mechanistic description of the molecular actors that drive state changes (**Extended Data Fig. 1b**) – have animated a conceptual framework for understanding cell fate choices for almost 60 years. Recent technological advances in single-cell epigenomic assays^7-12^ now provide the opportunity to imbue this allegorical landscape with molecular details by measuring both the overall shape of this epigenomic landscape, as well as the activity of master regulators, or "guy wires”, that influence cell fate decisions, within individual cells during normal tissue differentiation.

Hematopoietic differentiation serves as an ideal model for exploring the nature of multipotent cell fate decisions^13^. The hematopoietic system is maintained by the activity of a small number of self-renewing, long-lived hematopoietic stem cells (HSCs) capable of giving rise to the majority of blood cell lineages^13,14^ whereby multipotent cells transit multiple decision points while becoming increasingly lineage-restricted (**Fig. 1a**). The human hematopoietic system is an extensively characterized adult stem cell hierarchy with diverse cell types capable of phenotypic isolation with multi-parameter fluorescence activated cell sorting (FACS)^15^. This capacity for phenotypic isolation has enabled measurement of the epigenomic and transcriptional dynamics associated with sorted progenitors across human^16,17^ and mouse^18-20^ differentiation, providing a foundation for the dissection of regulatory variation in normal multi-lineage cellular differentiation.

**Figure 1.**
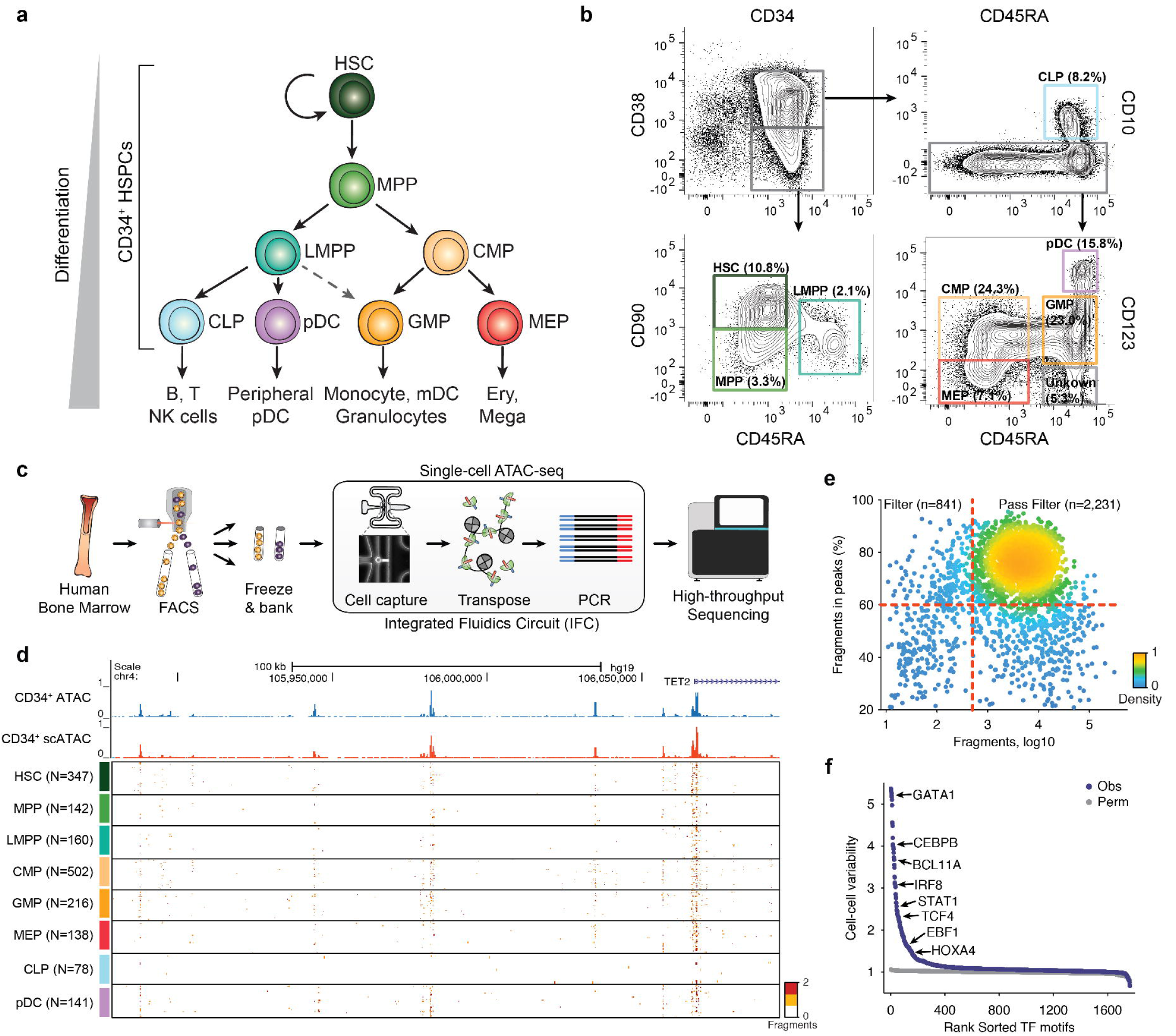
Single-cell ATAC-seq profiles chromatin accessibility within single hematopoietic progenitors. (a) A schematic of human hematopoietic differentiation. (b) Sorting strategy for CD34+ cells. (c) Single-cell ATAC-seq workflow used in this study. (d) Single-cell epigenomic profiles along the TET2 locus. (e) Fragments in peaks by library size plot show which cells pass filter; points are colored by density. (f) TF motif variability analysis in all single-cell epigenomic profiles collected for this study.

To define the single-cell epigenomic landscape of this developmental hierarchy, we used a single cell assay of transposase accessible chromatin by sequencing (scATAC-seq)^7^ to generate data from 10 sortable populations in human bone marrow or blood comprising multipotent and lineage restricted progenitors, which includes HSCs. We find that the regulatory landscape of human hematopoiesis is continuous, with cell surface markers reflecting epigenomic “basins” across this landscape. However, our single-cell analyses also uncovered substantial epigenomic heterogeneity within immunophenotypically-defined cellular populations, including epigenomic variability within multipotent progenitors strongly correlated along the dimensions of hematopoietic differentiation – evidence for regulatory lineage priming across multipotent progenitors. We also observe epigenomic variability within immunophenotypically-defined populations of common myeloid progenitor (CMP) and granulocyte-macrophage progenitor (GMP) cell types. Using RNA-seq, we confirm that GMPs are substantially heterogeneous on both epigenomic and transcriptomic levels, and describe a new strategy to enrich for subpopulations within GMPs at different developmental stages of a continuous myeloid differentiation trajectory. Last, we identify diverse regulatory dynamics at individual *cis*-regulatory elements, governed by relatively few overall *trans-factor* accessibility programs that shape the myeloid developmental trajectory. Thus, single-cell epigenomic analysis offers a platform for *de novo* discovery of cell types and states, defines regulatory variability within immunophenotypically pure populations, and captures the regulatory dynamics across a cell-resolved epigenomic landscape of differentiation first conceptualized by Waddington.

## Results

### Single-cell epigenomics of distinct hematopoietic cell types

We used FACS to isolate 8 distinct cellular populations from CD34^+^ human bone marrow, which included cell types spanning the myeloid, erythroid, and lymphoid lineages (**Fig. 1a,b**). In addition, we also profiled a CD34+CD38-CD45RA^+^CD123^-^ subset which has previously not been extensively functionally characterized^21^. We found that sorted cells (fresh) and cells cryopreserved after sorting (frozen) were highly comparable in their scATAC-seq data quality and yield (**Extended Data Fig. 1c-e**) and therefore performed all further scATAC-seq measurements on cells cryopreserved after FACS sorting (**Fig. 1c**). Together, this sorting strategy captures ~97% of all CD34+ cells (**Extended Data Fig. 1f**) and using post-sort analysis, we found that sorted cell types were on average 97% pure by cell surface marker immunophenotype^17^. Using this approach, we profiled the chromatin accessibility landscapes (referred to here as epigenomes for simplicity) across a total of 30 independent single-cell experiments representing 6 human donors, with each population assayed from two or more distinct donors. The data collected from each cell type roughly approximates the observed cell type frequencies from *in vivo* hematopoiesis (**Extended Data Fig. 1g**).

Aggregated single-cell profiles closely resemble bulk CD34+ ATAC-seq profiles (**Fig. 1d, Fig. Extended Data 1h,i**). Including previously published scATAC-seq data from LMPPs and monocytes^17^, this raw data set comprised 3,072 single-cell epigenomes across 32 integrated fluidic circuits (IFCs). Single-cell profiles were of consistent high-quality with 2,231 cells passing stringent quality filtering, yielding a median of 9,547 filtered fragments per cell with 76% of those fragments in peaks (**Fig. 1e; see methods**).

### Transcription factor activity inference using ChromVAR

We adapted our previous approach for finding cell-to-cell differences using transcription factor motifs to identify potential regulators of epigenomic variability^7^. In brief, this approach quantifies accessibility variation across single cells by aggregating fragments across accessible regions containing a specific transcription factor motif, then compares this aggregate accessibility to aggregate accessibility found in a set of background accessible regions matched for known technical confounders (GC content and overall accessibility). The method, we refer to as ChromVAR, provides a non-parametric measure of chromatin accessibility variation associated with transcription factor motifs, herein referred to as TF z-scores (Schep *et al*.). Using this approach we identify high-variance TF motifs representing known master regulators of hematopoiesis such as GATA1, BATF and CEBPB (**Fig. 1f**). In addition, hierarchical clustering of the TF z-scores generally classifies single cell profiles by their immunophenotypically-defined cell type identity (**Extended Data Fig. 2a**).

### Mapping single epigenomes on hematopoietic principal components

To map each single-cell to trajectories of hematopoietic development, we reasoned that co-variance of regulatory activity in bulk hematopoietic samples^22^ would provide a natural subspace for dimensionality reduction of our single-cell data. Therefore, we implemented a computational strategy that first identifies principal components (PCs) of variation in bulk ATAC-seq samples (**Extended Data Fig. 3a**), then scores each single-cell by the contribution of each PC. Cells are subsequently clustered using the Pearson correlation coefficients between these normalized PCs scores and all other cells (**Extended Data Fig. 3b**). For low-dimensional visual representation, we performed PCA on this correlation matrix (**Fig. 2a,b**), and display the first 3 principal components, which represent 93.7% of the variance (**Extended Data Fig. 3c**).

**Figure 2.**
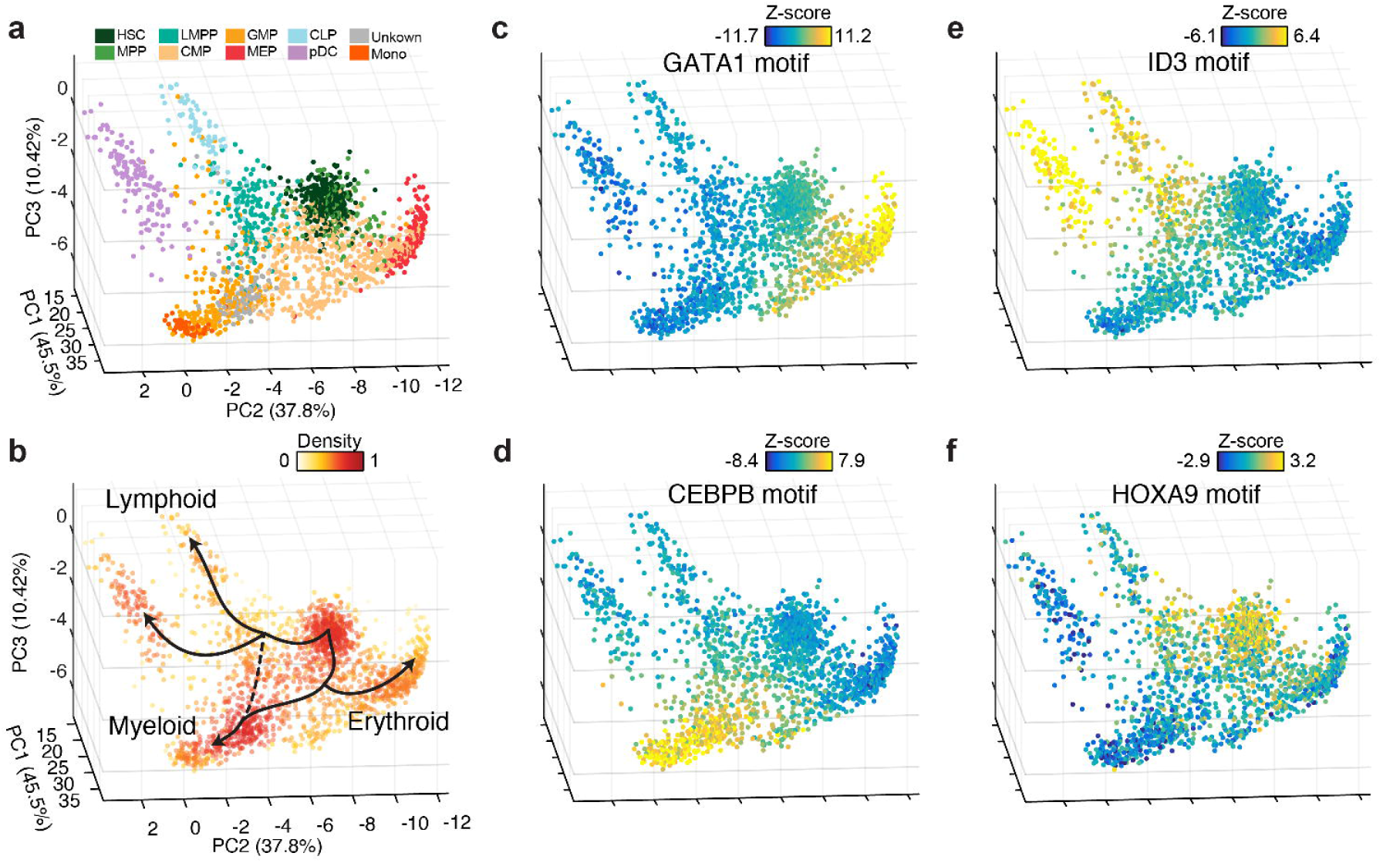
Lineage projection of human hematopoietic progenitors. Single-cell epigenomic landscape defined by PCA projection colored by (a) cell type identity using immunophenotype and (b) density (see methods) overlaid with nominal trajectories expected from the literature, as shown in Figure1a. PC projection colored by (c) GATA, (d) CEBPB, (e) ID3 and (f) HOXA9 motif accessibility TF z-score.

We validated this computational approach by down sampling bulk epigenomes across different patients to the approximate number of fragments per single-cell profile (**Extended Data Figure 3d**). We find that clustering samples down sampled to 10^4^ fragments using this PCA-projection approach closely follows sample clustering using the full data set^17^. We next down-sampled ensemble single-cell profiles to match the depths observed in single-cell experiments and quantified the expected mean error to be 1.95, 1.70 and 3.1% of the total signal for PCs 1-3 respectively (**Extended Data Fig. 3e,f**). To test our sensitivity for identifying intermediate cell states, we created synthetic mixtures from ensemble profiles, down-sampled to 10^4^ fragments and found these synthetic mixtures to closely follow the expected paths (**Extended Data Fig. 3i,j**). Thus this projection provides a descriptive landscape of human hematopoietic differentiation akin to Waddington’s conception (**Extended Data Fig. 1a,b**), while the layering of TF z-scores onto this representation provides insight into the "guy wires” that may underlie epigenomic changes during the differentiation process.

### Waddington landscape of human hematopoiesis: A manta ray

Projecting our scATAC-seq data using this method, the epigenomic landscape of human hematopoiesis resembles a swooping manta ray, with hematopoietic cell types generally localizing together (**Fig. 2a**). HSC (head, in dark green), LMPP and MPP (right and left shoulders, in teal and light green respectively), followed by broad groups of CMP and GMP cells (tan and light brown, respectively) that comprise the "body" of the figurative manta ray. Differentiation into CLP and pDC (right fins, blue and purple), MEP (left fin, red), and monocytes (tail, orange) appear as differentiation trajectories that swoop away from the body (**Fig. 2b**). Furthermore, motifs associated with master lineage regulators ID3, CEBPB and GATA1^13^ show continuous gradients of activity across lymphoid, myeloid, and erythroid development, while the HSC and LMPP compartment show higher accessibility associated with HOX motifs, previously shown to regulate stem cell activity^23,24^ (**Fig. 2c-f**). We also observe possible examples of cell type contamination from FACS misclassification, particularly between CMP:MPP, GMP:LMPP (separated by CD38) and GMP:CLP (separated by CD10) likely due to the continuous nature of the markers used to separate these subpopulations (**Extended Data Fig. 4**).

We found that CMP, GMP and MEP profiles appear markedly heterogeneous in this composite epigenomic space. We quantified the heterogeneity across each population by comparing the observed epigenomic variability in this space to down-sampled aggregate profiles and found CMPs to be the most heterogeneous cell type (p < 10^-111^), however, all cell types displayed statistically significant heterogeneity (**Extended Data Fig. 5a**). To further validate this ubiquitous epigenomic heterogeneity, we compared our results to results obtained after permuting peaks matched in mean accessibility and GC content (see methods) and find that the observed heterogeneity remains statistically significant with the exception of monocytes (p = 0.35; **Extended Data Fig. 5b,c**). Finally, single-cell TF z-scores (Schep *et al.*), which are calculated in an unrelated manner to this PCA projection, also exhibit significant variability (**Extended Data Fig. 5d**). Thus rather than identifying a series of discrete cellular states (**Fig. 1a**), these results suggest that the epigenomic landscape in early hematopoietic differentiation (HSC, MPP, LMPP and CMP) comprises a fairly broad basin of cellular states, while paths of later differentiation are more significantly canalized into distinct, however also continuous, myeloid, erythroid and lymphoid cell differentiation trajectories. We encourage the reader to explore the single cell epigenomic landscape using our interactive 3D viewer scHemeR (http://schemer.buenrostrolab.com/).

### *De novo* identification of uncharacterized epigenomic states

Given the observed limitations of our sort markers, we sought to define hematopoietic cell types *de novo* by applying k-medoids clustering on this principal component projection. We defined 14 unique clusters (see methods; **Fig. 3a,b**) that largely overlapped with previously-defined surface marker based definitions of human hematopoietic subsets, with several notable exceptions (**Fig. 3c**). We find that CMPs are separable into 4 distinct clusters, represented here as clusters 2-5. In addition, we also find that MEP, GMP and pDCs comprise two distinct clusters each, possibly representing early- and late-stage progenitor differentiation.

**Figure 3.**
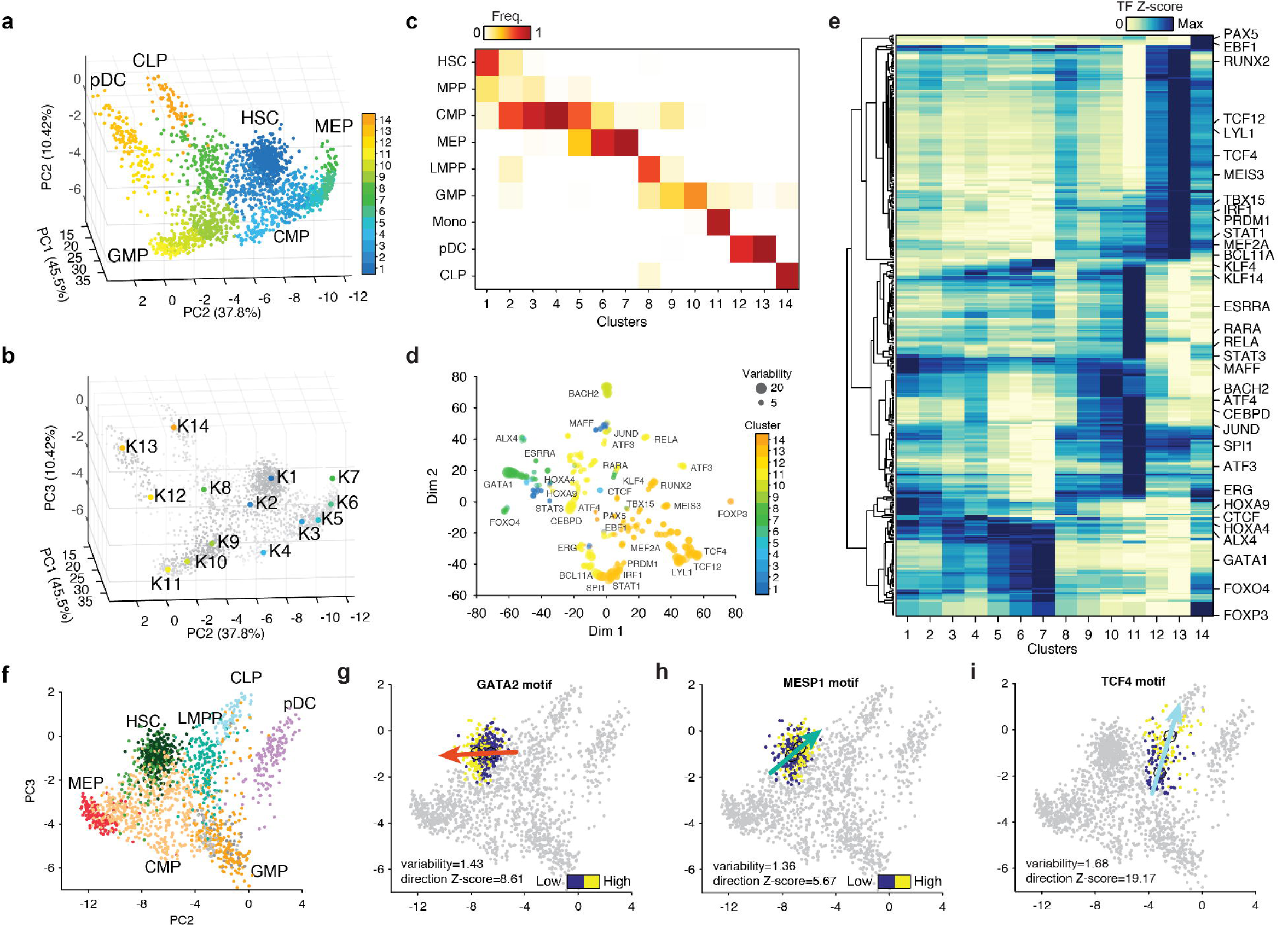
Molecular characterization of data-defined clusters. (a) Single cell epigenomic landscape defined by PCA projection, colored by data-driven cluster number. (b) Medoids of data-driven centroids depicted on the PCA sub-space. (c) Assignment matrix of data-driven clusters representing the percent frequency of immunophenotypically defined cell types. (d) t-SNE of TF activity z-scores colored by cluster with highest signal, points sized by cell-cell variability. (e) TF activity z-scores averaged across data defined clusters and hierarchically clustered, scores are normalized by the max value of each TF motif. (f) PC2 and PC3 projection of single-cell epigenomic profiles colored by the immunophenotypically defined clusters. (g-i) TF activity z-scores of HSC profiles for (f) GATA2 and (h) MESP1 motifs, or (i) LMPP profiles for TCF4 motif, arrows denote the direction of the signal bias and are colored by the target cell type.

We sought to further define these clusters by using transcription factor motif accessibility^7^. We visualized the activity patterns of TF motifs using t-SNE of the single-cell TF Z-scores (**Fig. 3d**) and hierarchical clustering of the mean scores across the 14 clusters (**Fig. 3e**). This analysis identified key hematopoietic regulators *de novo,* including motif associated with well-described master regulators such as GATA1 (erythroid), CEBPD (myeloid) and EBF1 (lymphoid) lineage-specifying factors^13^. Notably, we also find a specific HSC cluster of TFs that include HOX, ERG and MAFF motifs (see supplementary materials for a complete list of TF z-scores).

### Epigenomic variability within EIPP pure clusters

We next asked if we could see evidence of lineage-specific epigenomic signatures in stringently-defined progenitor subpopulations. To quantify this epigenomic variability, we created highly stringent cluster definitions that required classified cells to be both epigenomically (cluster-pure) and immunophenotypically pure (we term these EIPP clusters, see methods). We then computed TF z-scores (Schep *et al.*) for cells within each EIPP cluster (**Extended Data Fig. 6a**) and found substantial epigenomic heterogeneity within these subsets. We first explored the distribution of epigenomic variability observed in HSCs and found GATA2 and MESP1 TF z-scores to be significantly correlated to erythroid and lymphoid trajectories respectively (**Fig. 3f-h and Extended Data Fig. 6b-d**). Overall, we find much of the epigenomic variability within stringently defined EIPPs to be attributed to a developmental axis defined in our single cell PCA projection. However, we find CTCF, RELA and FLI1 motifs in HSCs to be significantly variable, however uncorrelated with any specific direction of differentiation, suggesting that these regulatory factors vary in a manner uncorrelated with the direction of hematopoietic development in this epigenomic landscape.

We next turned to LMPPs, a subset previously shown to demonstrate lineage priming toward dendritic (pDC), myeloid (GMP: Monocytes) and lymphoid (CLP:B-cell) fates in mice^25^. Interestingly, we found TCF4, STAT1 and CEBPE TF z-scores to be significantly correlated with directionality toward CLP, pDC and GMP differentiation, respectively (**Fig. 3i and Extended Data Fig. 6e-g**). Notably, TCF4 and CEBPE motifs were anti-correlated each with a unique direction toward CLP (lymphoid) and GMP (myeloid) differentiation, suggesting antagonism between myeloid/lymphoid differentiation programs, while the STAT1 motif appeared to be directed towards CLP (lymphoid) and pDC (dendritic) cell fates. Overall, we interpret this epigenomic variation associated with master regulators to be evidence of early lineage priming of progenitor cells, providing epigenomic support for previous reports showing strong lineage bias within otherwise pure subsets^25,26^.

### Cell state transitions defining myeloid cell differentiation

Recent findings have identified significant functional and transcriptional heterogeneity within the mouse^27,28^ and human^29^ CMP progenitor cells, and suggest that CMPs can be further partitioned into myeloid and erythroid committed progenitors. In light of these new discoveries, we reasoned that our data may provide insight into the regulatory components defining myeloid/erythroid specification, previously confounded in bulk epigenomic studies due to the apparent heterogeneity within CMP cells. In concordance with these findings we find four CMP clusters, with significant variability across GATA1, BCL11A and SPI1 (PU.1), TFs implicated in myeloid/erythroid specification (**Fig. 4a**). Furthermore, we identified 1,801 differentially accessible regions within CMPs and using k-medoids clustering, identified distinct regulatory patterns across the four CMP subsets (see methods; **Fig. 4b**). Importantly, the set of differential enhancers enriched in CMP clusters K3-K5 included two previously validated enhancers^30^ shown to be important regulators of the master erythroid factor GATA1 (**Fig. 4b**). Together with the TF z-scores we assigned CMP clusters as CMP-K3 (early erythroid), CMP-K5 (late erythroid), CMP-K4 (unknown) and CMP-K2 (myeloid primed).

**Figure 4.**
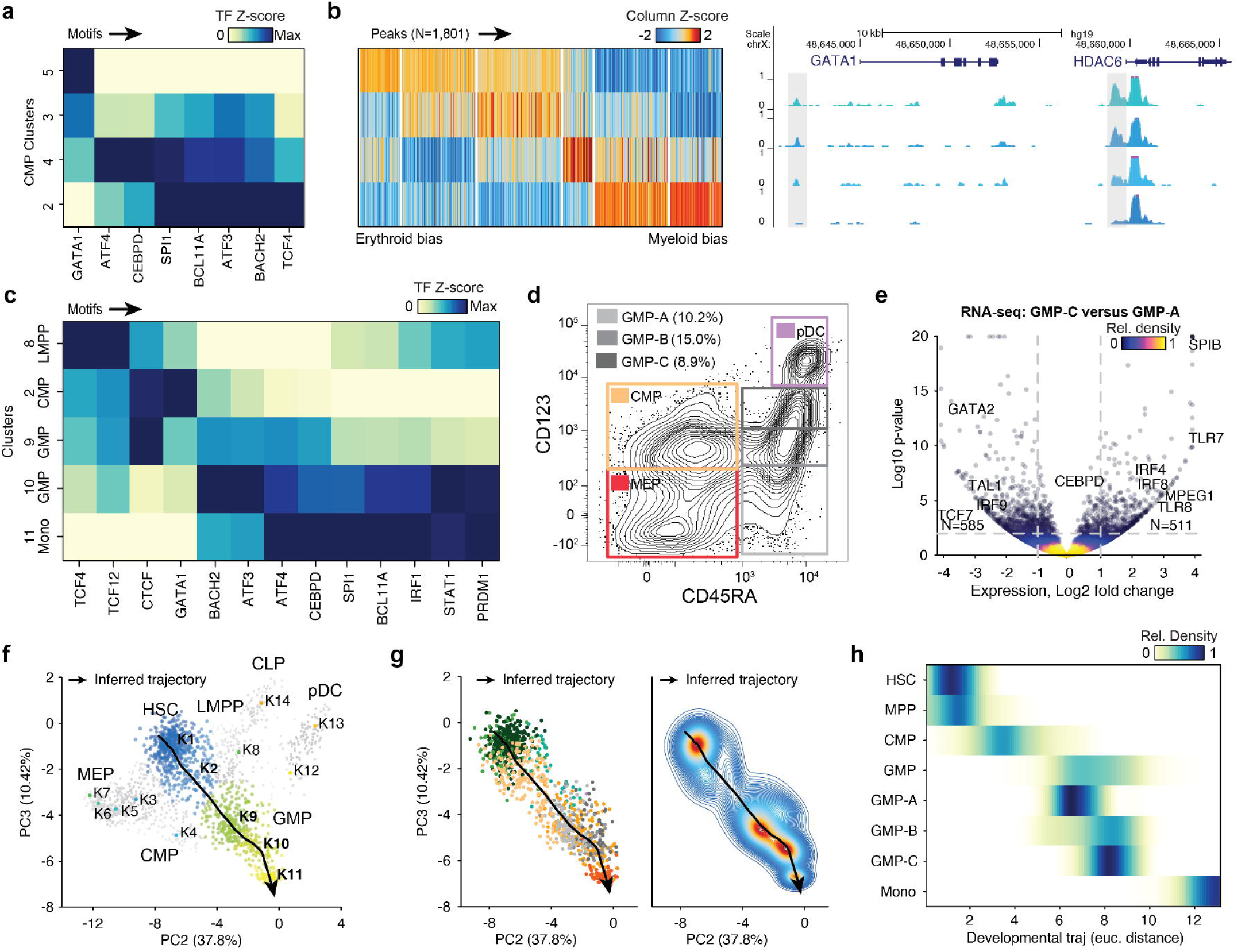
Identifying the continuous myeloid differentiation trajectory. (a) Differential motifs and (b) putative enhancers across CMP clusters (K2-5). Motifs are normalized by max-min values and (left) enhancers are normalized as z-scores and clustered using k-medoids. (Right) accessibility at GATA1 locus across the CMP clusters highlighting (grey) two validated^30^ enhancers of GATA1. (c) TF z-scores across clusters for TF motifs significantly differential across clusters K9 and K10, scores are normalized by the max value of each TF motif. (d) Sorting schema for different GMP progenitors defined by CD123 expression, marked by CD123 low (GMP-A, light-grey), CD123 medium (GMP-B, grey) and CD123 high (GMP-C, dark-grey). (e) RNA-seq log_2_-fold-change and-(log p-value) for expressed genes comparing GMP-C and GMP-A. (f) PC2 by PC3 projection of single-cells highlighting cells progressing through the inferred myeloid developmental trajectory (black line), colored and labeled by the data driven clusters, all other cells are shown in grey. (g) Single cells used for the myeloid trajectory colored by (left) their immunophenotypic sort identity (colors as in Fig. 1) or (right) their density along the trajectory. (h) Density of myeloid progression scores for immunophenotypically defined cell types, including the GMP subsets.

In addition to the heterogeneity within CMP cells, the GMP clusters K9 and K10 show significant differences in accessibility among myeloid-defining factors such as CEBP and SPI1 (PU.1) (**Fig. 4c and Extended Data Fig. 7a**). In an effort to further define molecular differences within immunophenotypically defined GMPs, we sought to identify cell surface markers that may differentially enrich for K9 and K10 GMPs. We hypothesized that CD123 expression may correlate with early and late GMP differentiation for two reasons: (i) the UNK population, which is CD123^-^, is enriched in the GMP K9 cluster and (ii) CD123, also known as IL3R, is a receptor for the myeloid promoting cytokine IL3. We therefore performed scATAC-seq, bulk ATAC-seq^31,32^ and bulk RNA-seq on cells from each of three distinct bins of CD123 expression (**Fig. 4d**). Projection of the three cell fractions using scATAC-seq data identified GMP-A as enriched in GMP K9, while GMP-B and GMP-C are enriched for GMP K10 (**Extended Data Fig. 7b,c**). In addition, bulk ATAC-seq and RNA-seq data revealed significant epigenomic and transcriptomic differences across the GMP-A and GMP-C populations (**Fig. 4e and Extended Data Fig. 7d-f**). The list of differentially expressed genes included important developmental regulators, including down regulation of HSPC TFs GATA2 and TAL1 and up regulation of myeloid genes SPI1 (PU.1), IRF8, TLR7 and MPEG1 in the GMP-C cell population (**Fig. 4e**). Altogether, this single-cell and bulk analysis supports our data driven approach for uncovering developmental differences in human hematopoiesis and suggests that CD123 expression marks molecularly distinct early and late GMP progenitor populations.

### Regulatory dynamics along myeloid cell differentiation

Spurred by this strong epigenomic signal of population-level heterogeneity along myeloid differentiation, we sought to project single-cells onto an inferred developmental trajectory. Integrating both recently published work and our data described above, we reasoned that myeloid differentiation proceeds through: HSC (K1), the myeloid biased CMP (K2), GMP-A (K9), GMP-C (K10) and monocytes (K11) (**Fig. 4f**). Notably, this myeloid trajectory also represents the shortest path from HSCs to monocytes in our projected hematopoietic landscape. We therefore assigned cells within these clusters to a continuous myeloid developmental order (see methods; **Fig. 4f-h and Extended Data Fig. 7g**).

We next characterized TF dynamics across myeloid development by mapping TF z-scores to cells along this continuous myeloid differentiation trajectory (**Extended Data Fig. 8**). Using this approach, we find 6 clusters of TF activity profiles that occur during myeloid development (**Fig. 5a**). TF motifs associated with regulators HOXB8 and GATA1 motifs (cluster 1) are active in HSCs and are lost upon differentiation to CMPs. Interestingly, loss of GATA1 motif activity begins within the HSC compartment, while HOXB8 activity is lost at the transition of HSC to CMP differentiation further demonstrating that GATA motif accessibility may be involved in lineage priming of HSCs (**Fig. 5b**). We also find two distinct modes of activation of myeloid-associated TFs; cluster 4 TFs (CEBP and SPIB) display early and gradual gain in activity beginning within CMPs, while cluster 5 TFs (STAT1, IRF8 and BCL11A) increase sharply in activity across the GMP-A to GMP-C transition, implicating CEBP motif activity as the initiating factor for myeloid-erythroid specification (**Fig. 5c**). In addition to the activity patterns associated with canonical myeloid-defining factors, we also identify a pulse of activity within CMPs from cluster 2 TFs (TCF3/12, TFs up regulated in CLP/pDC), which may reflect transient activation of a lymphoid program within the pre-committed myeloid-biased CMPs. Overall, resolution of the order of these dynamic trans-*factors* has been previously obscured in bulk epigenomics studies, in part, due to the cellular heterogeneity within CMPs. Here, single-cell epigenomics resolves the temporal dynamics of master regulators in myeloid cell development.

**Figure 5.**
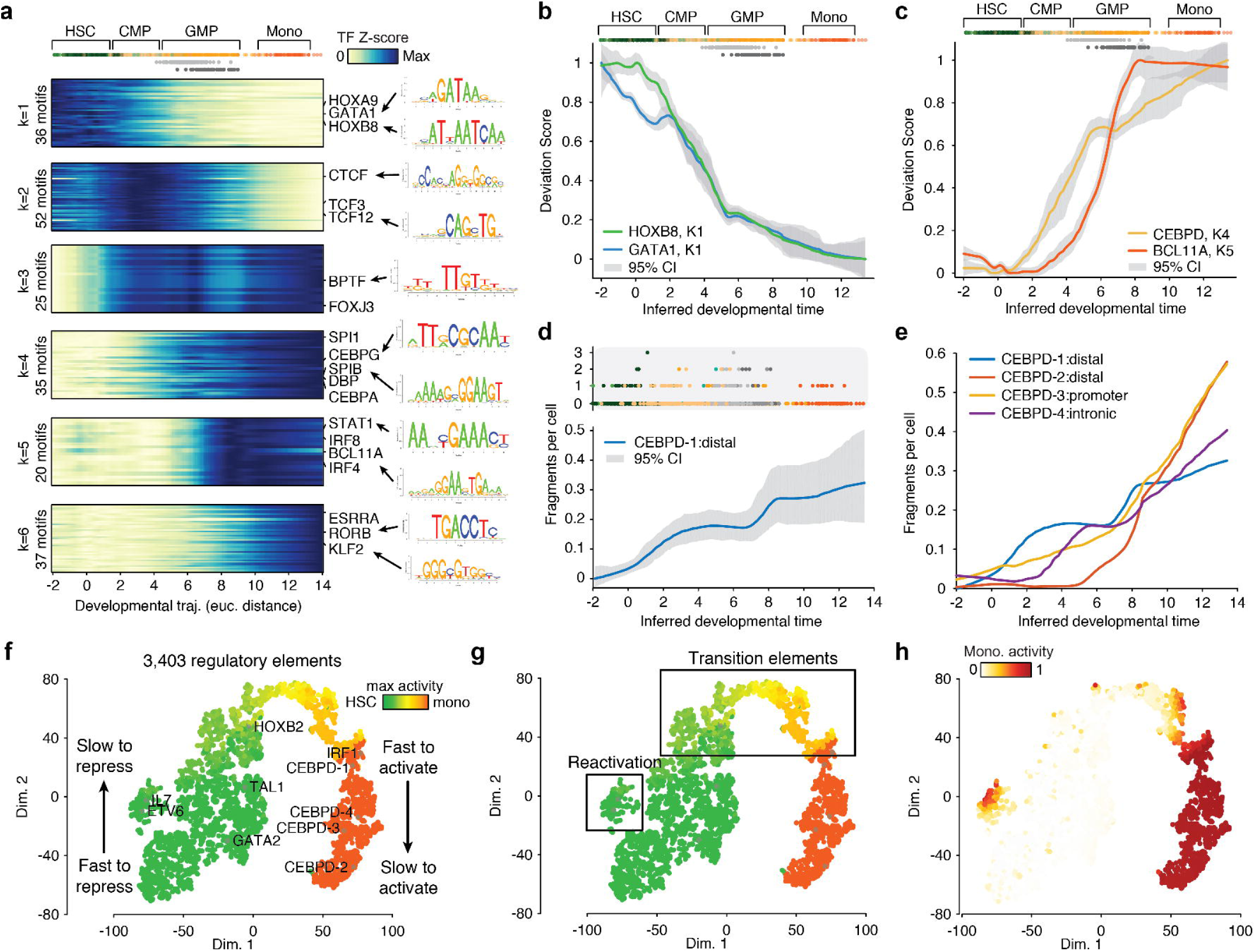
Regulatory dynamics across a continuous myeloid differentiation trajectory. (a) K-medoids clustering of TF activity (left) and PWM logos (right) for dynamic TF motif profiles across myeloid development. (b,c) Smoothed profiles of motif TF z-scores in myeloid progression for (b) HSC active TFs GATA1 (blue) and HOXB8 (green), and (c) monocyte active regulators CEBPD (yellow) and BCL11A (red), error bars (grey) denote 95% confidence intervals. (d) (top) Cells are colored by their immunophenotype and ordered by their progression score. (bottom) Smoothed *cis*-regulatory activity profile (fragments per cell) of a CEBPD distal element across myeloid development, error bars (grey) denotes 95% confidence intervals. (e) *Cis*-regulatory dynamics across four regulatory elements near the myeloid regulator CEBPD. (f,g) t-SNE embedding of *cis*-regulatory profiles colored by the maximum activity along the developmental axis. Plots highlight (f) slow or fast activation or repression and (g) reactivation or transition elements. (h) Activity of each *cis*-regulatory element in monocytes projected into the t-SNE embedding shown in panels (f,g).

We next sought to identify *cis*-regulatory activity dynamics underlying myeloid differentiation. To do this, we filtered for regulatory elements with high fragment counts and with significant variability across the ordered cells (see methods), identifying 3,403 *cis*-regulatory elements for analysis. From this list of regulatory elements we found highly heterogeneous patterns of activity characterized by their activation timing (**Extended Data Fig. 9a-d**). For example, within the regulatory elements surrounding the myeloid regulator CEBPD (numbered for simplicity), the distal element CEBPD-1 was “fast-to-activate” and showed step-wise gains of activity while the distal element CEBPD-2 was “slow-to-activate” and showed a discrete pulse of activity (**Fig. 5d,e**).

To visualize the complete repertoire of dynamic regulatory profiles, we used t-SNE to order elements based on their accessibility changes over this trajectory (see methods). This projection reveals multiple axes of regulatory element behaviors, ranging from fast- to slow-to-repress HSC regulatory elements and fast- to slow-to-activate monocyte regulatory elements (**Fig. 5f**). We also observe a collection of “transition” *cis*-regulatory elements that exhibit peak accessibility at intermediate stages of myeloid development, as well as “reactivation” elements that initially are lost and subsequently reactivated in later stages of myeloid differentiation (**Fig. 5g,h**). Altogether this single-cell resolved regulatory analysis of myeloid cell development revealed 6 discrete clusters of TF motif activity, however, we find that these relatively simple patterns of *trans-dynamics* yields highly diverse *cis*-regulatory profiles likely arising from the large combinatorial control of trans-factors on their cognate regulatory elements (**Extended Data Fig. 9e**).

## Discussion

We used single-cell epigenomic analysis to identify regulatory heterogeneity and continuous differentiation trajectories in a human differentiation hierarchy. We find that immunophenotypically-defined cell populations often flow from one state to another within the epigenomic landscape, in support of a more continuous model of hematopoietic cell differentiation. This continuous nature can be seen in i) our single-cell epigenomic projection, ii) epigenomic bias consistent with lineage priming within EIPP clusters and iii) asynchronous yet step-wise gains of accessibility at genomic loci during myeloid development. Furthermore, single cell epigenomes can be aggregated to define, in an immunophenotype-agnostic and data driven manner, unique *cis*-regulatory elements active at different stages of the developmental trajectory or in different cell states. The intersection of phenotypically-associated variants with these regulatory elements may provide further insight into cell types and differentiation stages relevant to specific disease etiologies^17,33,34^.

We have produced an intuitive representation of the density of hematopoietic precursor states as a function of epigenomic “similarity” that is reminiscent of Waddington’s conceptual landscape of differentiation. However, it remains uncertain to what degree this single-cell epigenomic projection is interpretable in terms of transition kinetics and transition state potentials, as is expected by the most physical interpretation of this landscape concept. Indeed, none of the dynamic aspects of state changes are explicitly measured in our data set. Thus, we are unable to distinguish, for example, whether HSCs are relatively stable populations of primed cells or if they dynamically fluctuate in their observed “priming” over time. A more kinetically quantitative instantiation of the epigenomic landscape will likely require direct estimates of transition probabilities, either through sorting marker-pure HSPC subpopulations followed by time-point analysis of differentiation or through the emerging repertoire of CRISPR-based tools for kinetic lineage tracking^35,36^, which would unravel the dynamic nature of chromatin state changes and the potential for reversibility.

We anticipate future work will experimentally or computationally pair epigenomic and transcriptomic data to enable an integrative regulatory model of dynamic cellular processes, providing insight into the temporal function of *trans* regulators and *cis*-regulatory elements. However, it remains to be seen to what degree transcriptomic and epigenomic measures of cellular phenotypic states are correlated, given disparate timescales for gene expression, chromatin remodeling, and RNA degradation. In addition, one might anticipate epigenomic memory-encoded processes, such as lineage priming, to be strongly represented in single-cell epigenomic profiles along developmental trajectories. Indeed, pervasive lineage priming in mouse hematopoietic progenitors has been observed^25,26^, and recent work has shown mouse lineage-primed HSCs to exhibit extensive epigenomic differences (assayed using ATAC-seq) with little to no transcriptomic differences^37^. However, one might expect other biological phenomena that require fast stimulus-response programs to bypass chromatin changes, and exhibit differences in RNA or protein modifications^38^. In either case, we anticipate that single-cell epigenomics assays will provide a platform for unraveling *cis-*and *trans-*regulatory changes associated with dynamic differentiation trajectories or tissue identity in an immunophenotypically unbiased and data driven manner.

---

**Extended Data Figure 1.**
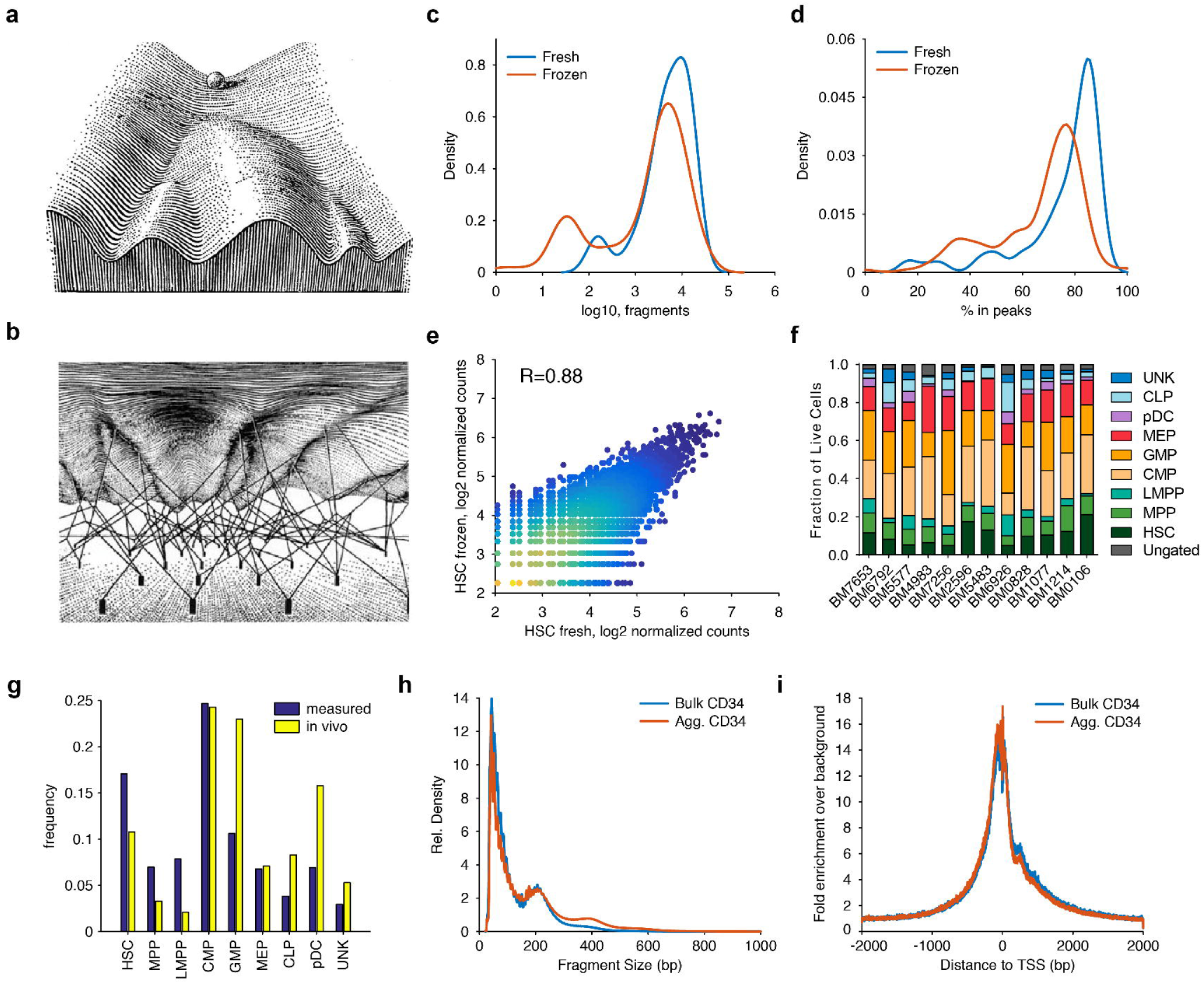
Quality characteristics of single-cell epigenomes. (a,b) Waddington landscape representing (a) the sinuous epigenetic landscape wherein a cell (ball) can roll down different cell fates, and (b) "guy-wires" that shape the epigenetic landscape. (c,d) Comparison of scATAC-seq from ‘fresh’ (blue) and cells frozen after FACS sorting (red). Profiles from HSCs showing the (c) fragment yield per cell and (d) fraction of fragments in peaks. (e) Comparison of the log_2_ accessibility between donor-matched fresh and frozen aggregate accessibility profiles, R=0.88. (f) Fraction of immunophenotypically defined cell types from CD34^+^ cells for each bone marrow donor, ungated cells are marked in grey. (g) The measured (blue) and average *in vivo* (yellow) frequency of cells in the data set. (h) Fragment size (bp) distribution and (i) enrichment at transcription start sites (TSSs) for bulk (blue) and aggregate single-cell (red) profiles.

**Extended Data Figure 2.**
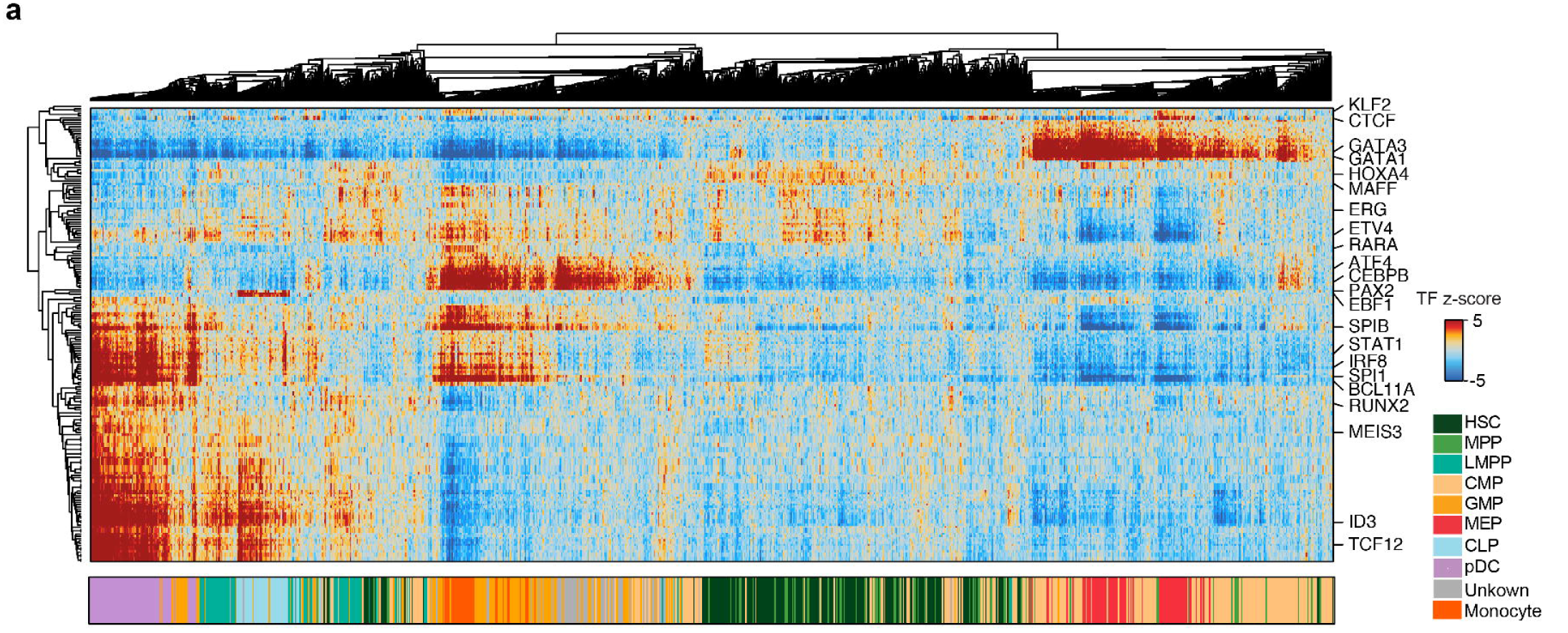
Hierarchical clustering of epigenomic profiles. (a) (top) Hierarchical clustering of single-cell epigenomic profiles (columns) and TF z-scores (rows), (bottom) single cell profiles colored by their sorted immunophenotype identity.

**Extended Data Figure 3.**
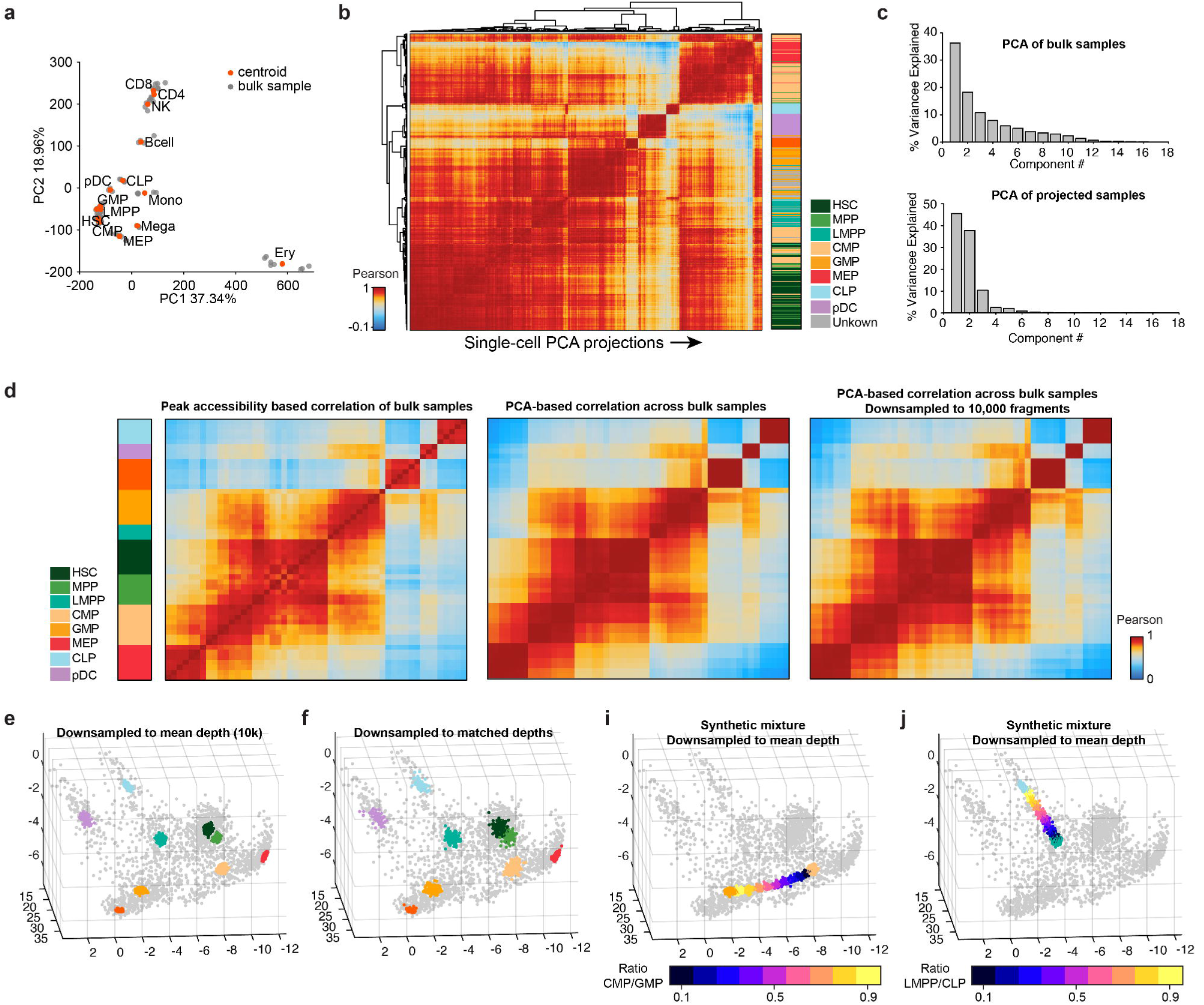
Development of an analytical framework for PCA projection. (a) PC1 and PC2 from PCA of fragments in peaks for bulk samples (grey) and their centroids (red). (b) (left) Hierarchical clustering of single-cell epigenomic profiles scored by bulk PCs, (right) profiles colored by their sorted immunophenotype identity. (c) Percent variance explained for each PC from (top) PCA derived from bulk data using fragments in peaks and (bottom) PCA of the PCA projected subspace (see methods). (d) Bulk sample-sample correlation across (left) fragments in peaks, (middle) PCA projection values and (right) PCA projection values after down sampling data to 10,000 fragments per sample. Far left represents the sorted immunophenotype of each bulk sample. (e,f) Mean of immunophenotypically defined single cell profiles down sampled to (e) 10,000 fragments or (f) matched fragment counts to the observed single-cell data set. (i,j) Synthetic mixtures of two immunophenotypically defined single-cell profiles down sampled to 10,000 fragments with mixtures of (i) CMP/GMP and (j) LMPP/CLP cell types, unmixed cell types from panel (e) are shown for reference.

**Extended Data Figure 4.**
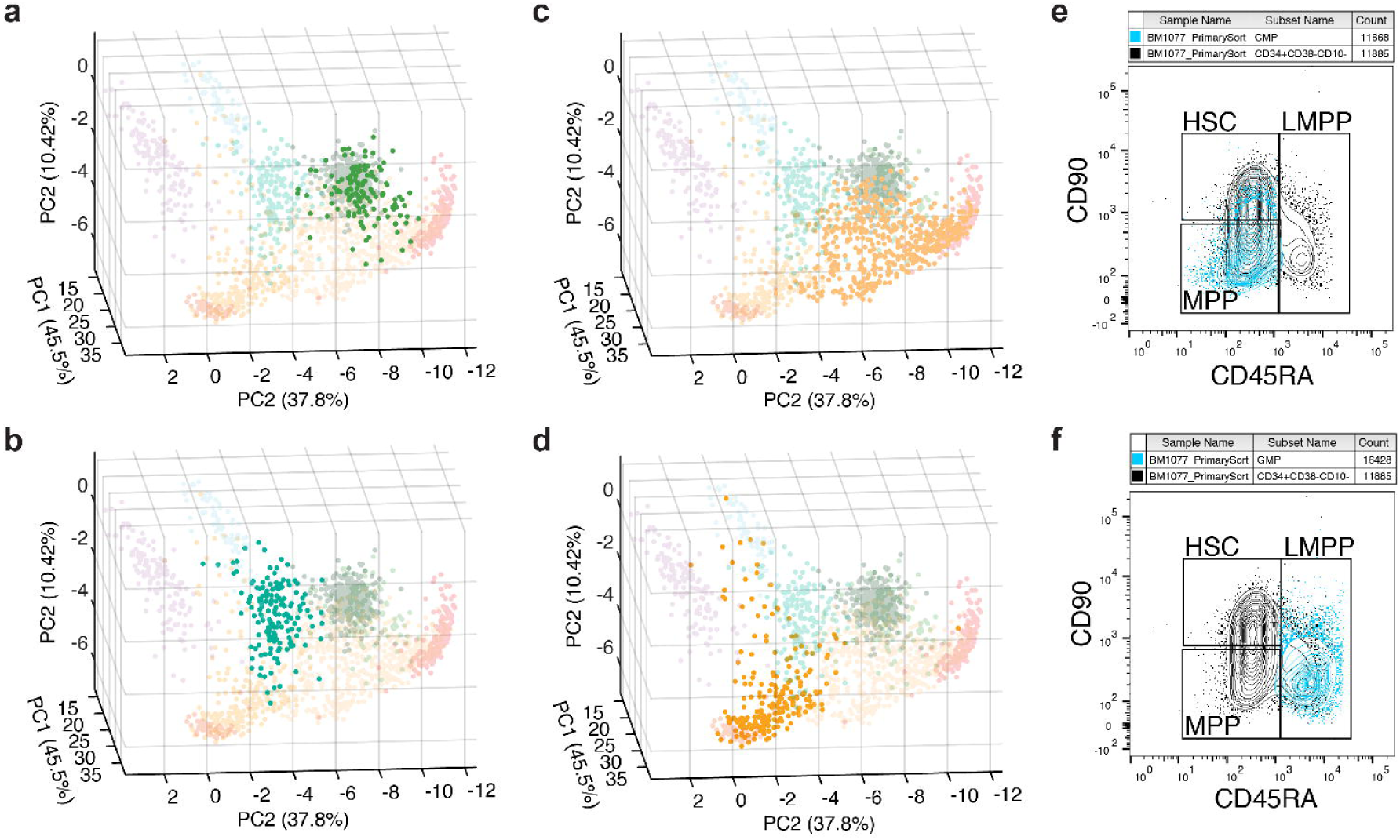
Sources of error in FACs sorting. (a-d) PCA projection of highlighted cell types for (a) MPP, (b) LMPP, (c) CMP and (d) GMP. (e,f) Flow cytometry back gating of (e) CMPs and (f) GMPs to show that a subset of cells exhibit CD90 and CD45RA cell surface marker expression without the CD38 signal. These potentially miss-gated CMPs localize to the MPP gate while miss-gated GMPs localize to the LMPP gate.

**Extended Data Figure 5.**
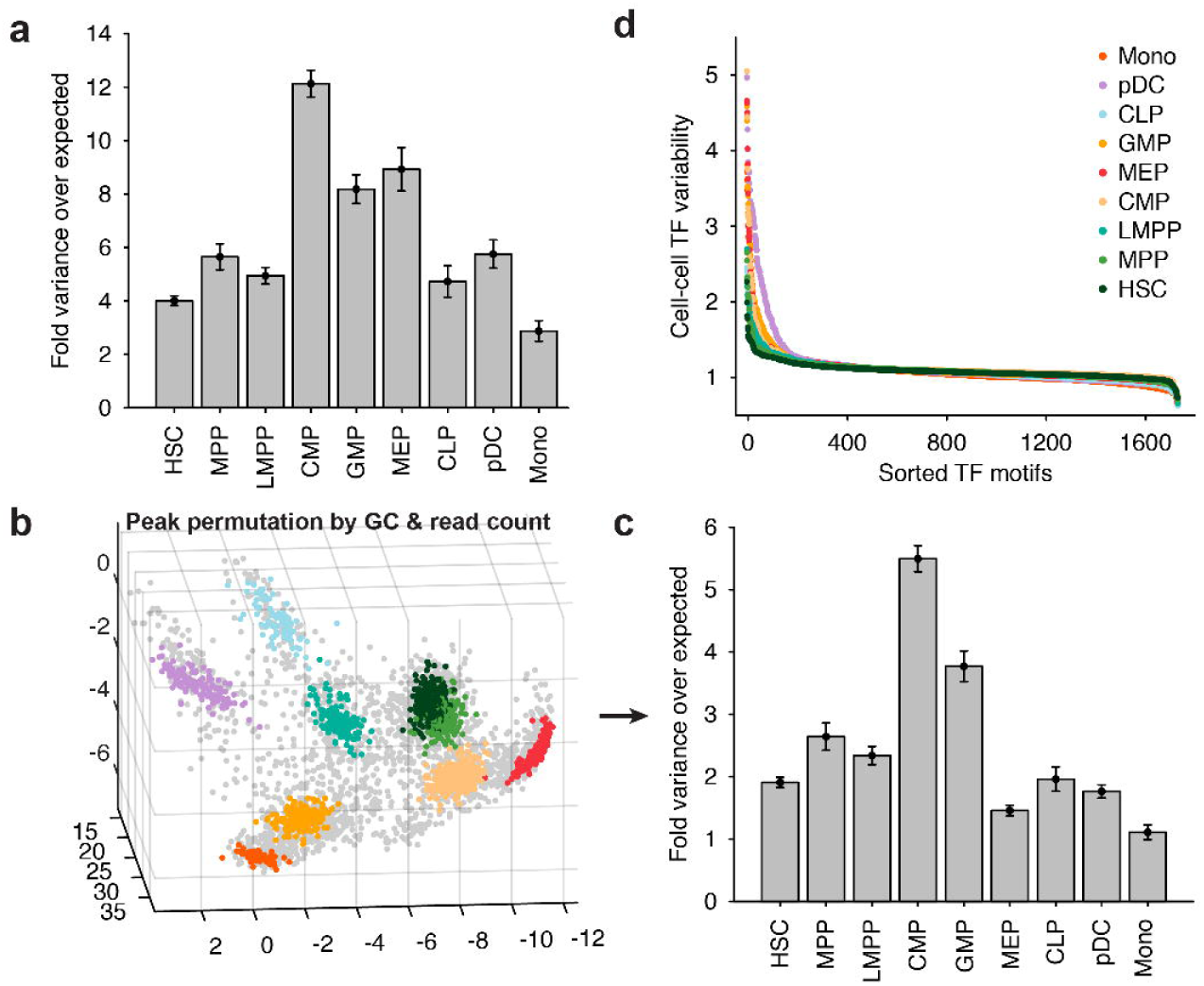
Tests for determining significant epigenomic variability within cell types. (a) Fold variance of the PCA projection over the variability due to count noise, determined by down sampling each cell type to matched sequencing depths, for each cell type. Error bars represent 1 standard deviation, and are estimated using bootstrap sampling (1,000 iterations) of cells. (b) Peaks permuted by their GC content and mean fragment count for each aggregate single-cell profile, then projected onto the PC subspace. (c) Fold variance over expected for each cell type, as shown in panel (a) using the permuted scores shown in (b). (d) TF variability sorted by the rank score for each cell type.

**Extended Data Figure 6.**
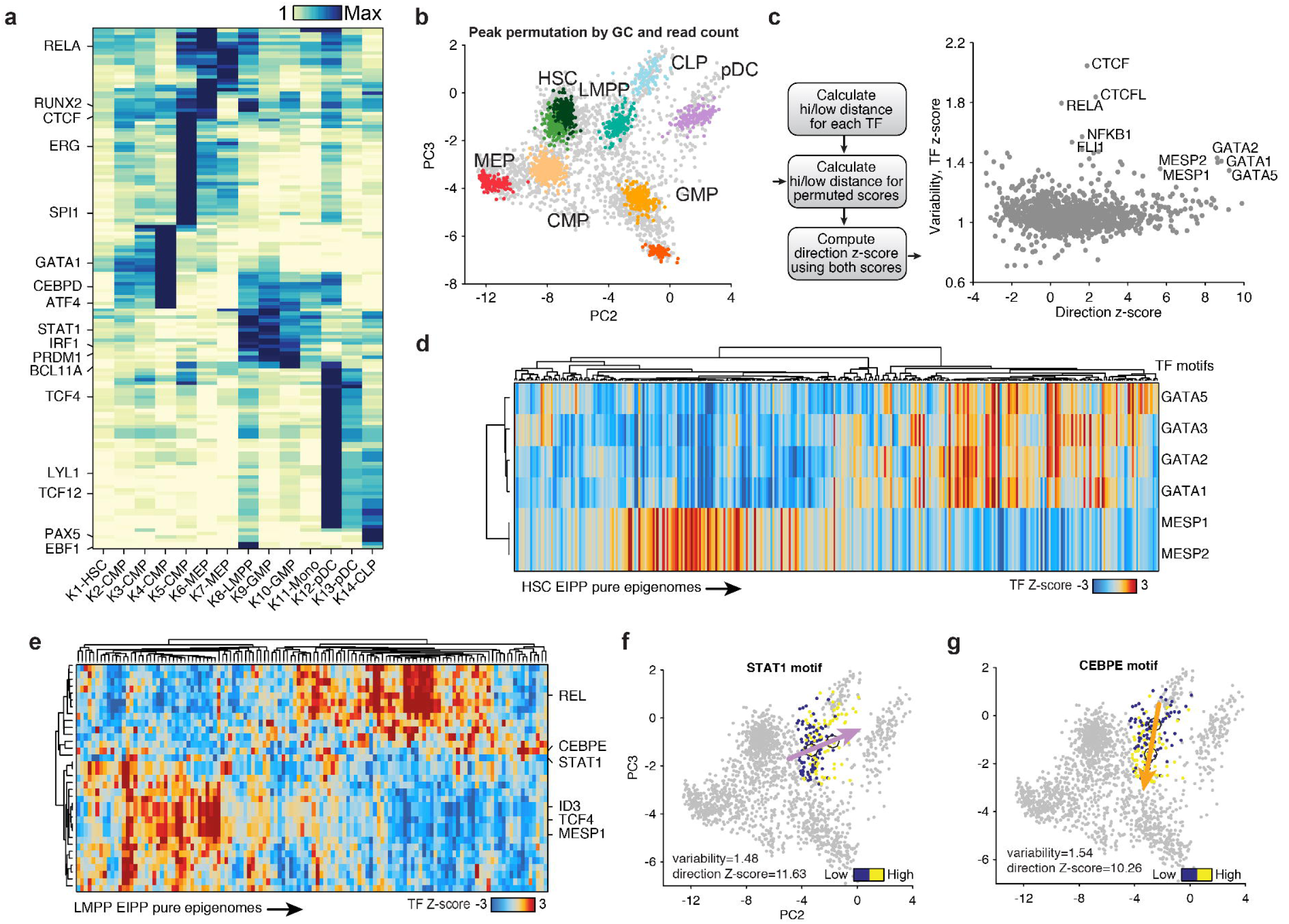
Epigenomic variability within defined cell states. (a) Cell-cell variability across TF motifs within each EIPP cluster (see methods). (b) Peaks were permuted by their GC content and mean fragment count for each aggregate single-cell profile, single cell profiles were then projected onto the PC subspace. (c) (left) Schematic for determining direction z-score using permuted PCA scores (n=50) described in plane (b), (right) TF variability and direction z-score for each TF motif within the HSC EIPP cluster. (d,e) Hierarchical clustering of single-cell (d) HSC and (e) LMPP EIPP profiles (columns) by TF motifs appearing as highly variable and directional (rows). (f,g) PC2 and PC3 projection of single LMPP profiles colored by high (yellow) or low (blue) TF activity z-scores for (f) STAT1 and (g) CEBPE motifs, arrows denote the direction of the signal bias.

**Extended Data Figure 7.**
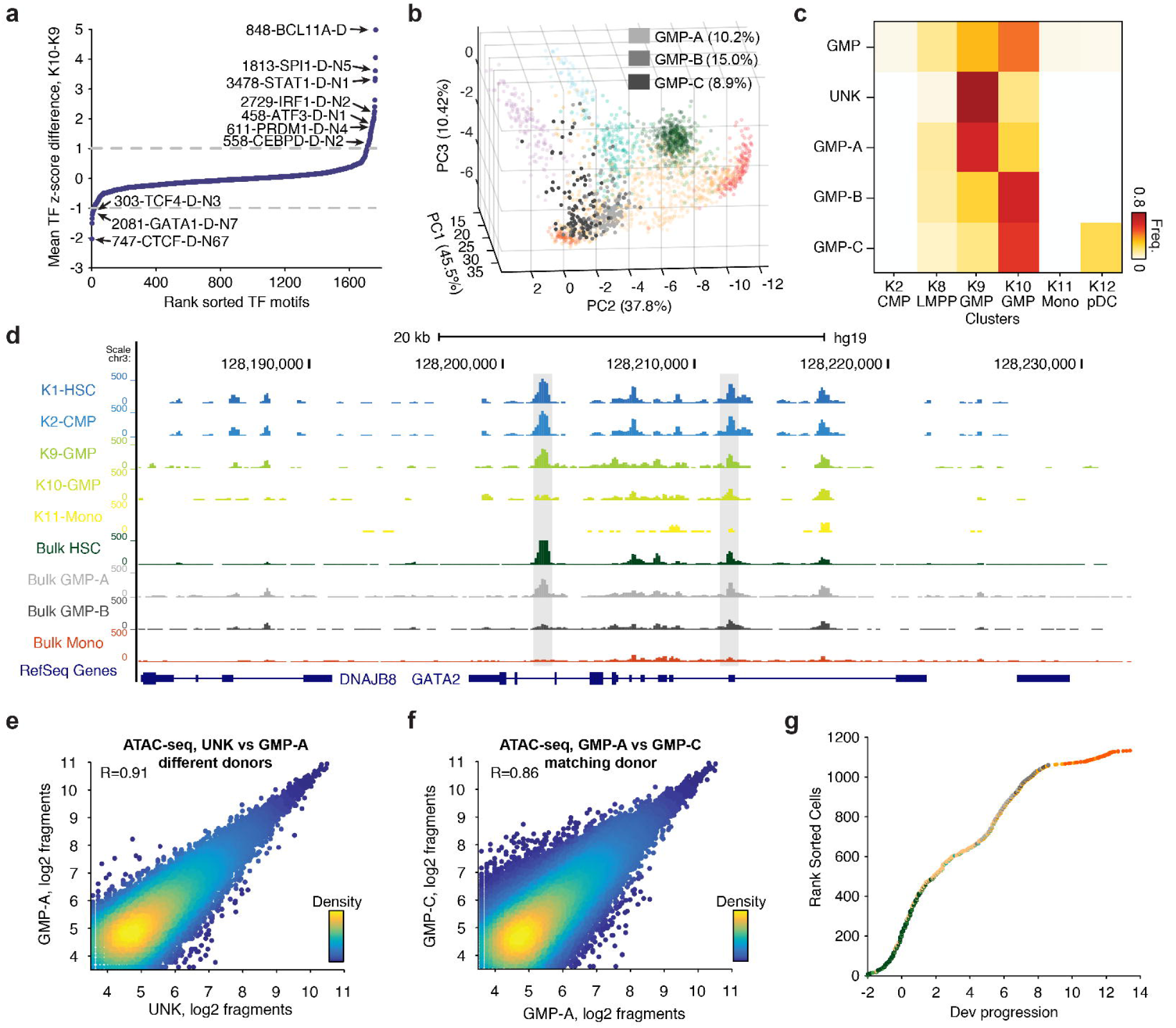
Molecular heterogeneity within GMPs. (a) TF motifs sorted by their rank difference in the mean TF z-score between GMP K10 and K9 clusters, important regulators are highlighted. (b) PCA projection of (light grey) GMP-A, (grey) GMP-B and (dark grey) GMP-C single-cell profiles. (c) Frequency of single-cell profiles from differing GMP sorted immunophenotypes within previously defined epigenomic clusters. (d) Genome browser track highlighting two differentially accessible regions surrounding the GATA2 gene. (e-f) Scatter plots colored by density denoting log2 fragments in individual peaks across samples for (e) UNK versus GMP-A and (f) GMP-A versus GMP-C profiles. (g) Single-cell profiles colored by their immunophenotypic origin rank sorted by myeloid developmental progression.

**Extended Data Figure 8.**
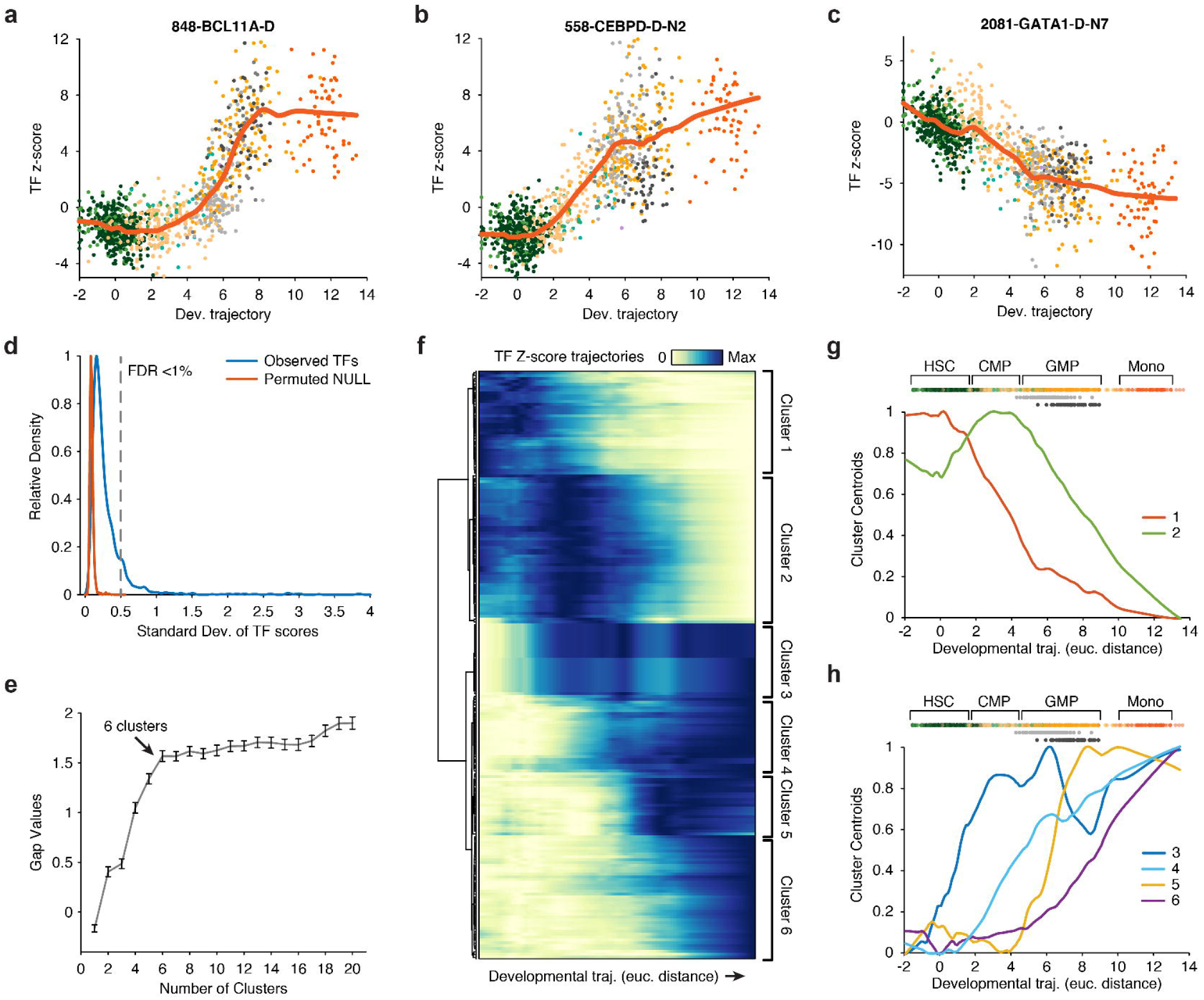
Transcription factor dynamics across myeloid cell differentiation. (a-c) Single-cell myeloid progression and TF z-scores for motifs (a) BCL11A, (b) CEBPD and (c) GATA1, smoothed motif activity trajectories are shown in red. (d) The standard deviation of smoothed TF activity scores, as shown above, for observed scores (blue) and cells randomly permuted (red), dotted line represents an FDR greater than 1%. (e) Gap values for 1 to 20 k-medoids clusters, the optimal cluster number 6 is highlighted. (f) Hierarchical clustering for smooth TF z-score trajectories (n=202 motifs), cluster identified using k-medoids are highlighted. (g,h) Centroids for each k-medoid TF cluster for (g) early and (h) late activity TFs, cells ordered by myeloid development (top) are shown for reference.

**Extended Data Figure 9.**
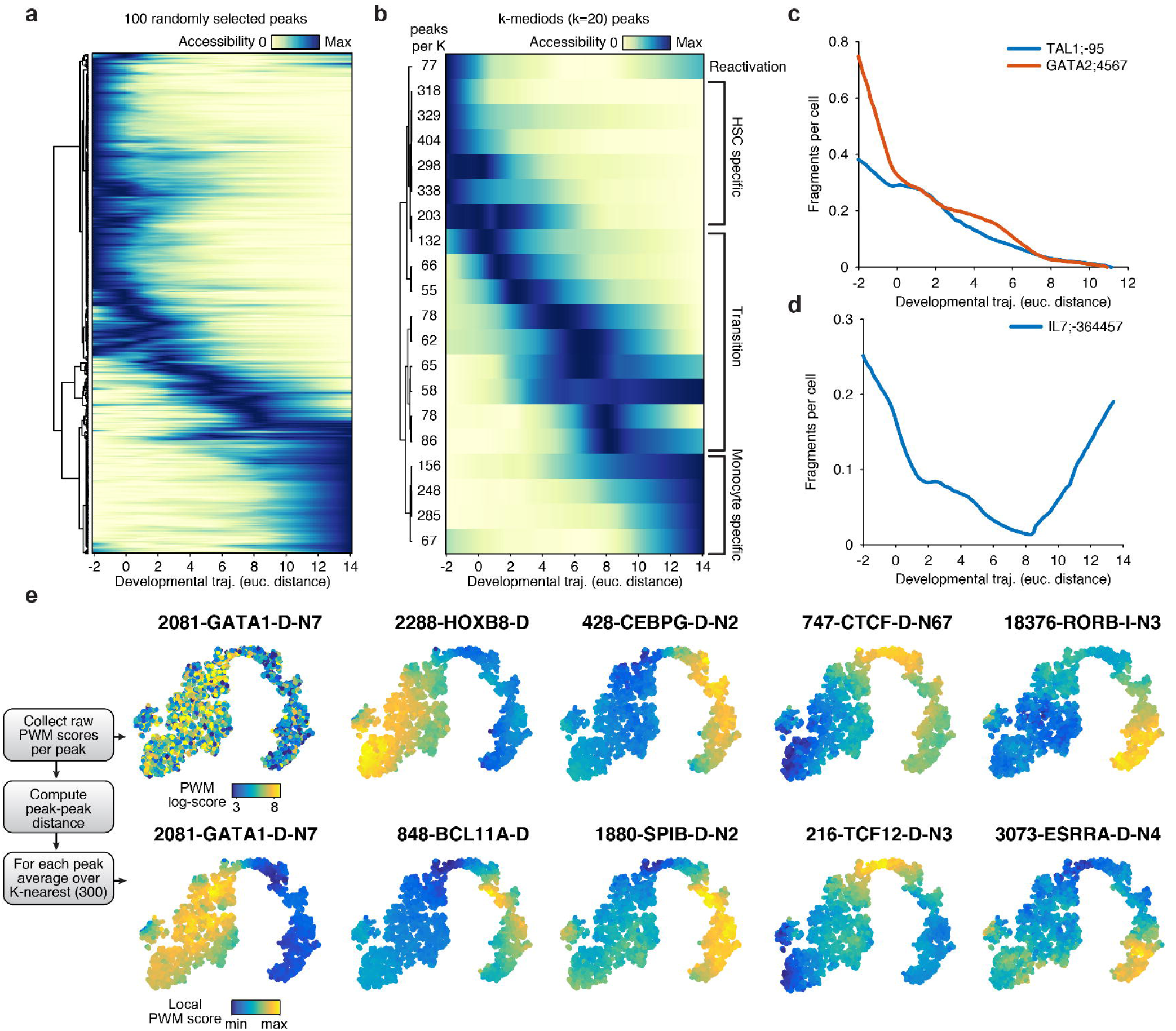
TF dependent *cis*-regulatory element dynamics. (a) Hierarchical clustering of 100 randomly selected accessible elements smoothed as a function of myeloid progression. (b) Hierarchical clustering of k-medoids centroids across all peaks, values (left) denote number of peaks per cluster and labels (right) denote categorization into different global patterns. (c,d) Example *cis*-regulatory dynamics across myeloid development for (c) HSC active peaks and (d) a reactivated peak. (e) (left) Schematic for determining motif enrichment across similar peak profiles, (right) raw log PWM score or averaged local PWM score for a representative subset of dynamic TF motifs across myeloid development for the t-SNE embedding shown in Fig. 5.

## Acknowledgements

This work was supported by National Institutes of Health (NIH) P50HG007735 (to H.Y.C. and W.J.G.), Stinehart-Reed Foundation (R.M. and H.Y.C.), Life Extension Foundation (H.Y.C.), U19AI057266 (to W.J.G.) and the Rita Allen Foundation (to W.J.G.) and the Baxter Foundation Faculty Scholar Grant and the Human Frontiers Science Program (to W.J.G). J.D.B. acknowledges support from the Harvard Society of Fellows and Broad Institute Fellowship. R.M. is a New York Stem Cell Foundation Robertson Investigator and Leukemia and Lymphoma Society Scholar. We thank members of Greenleaf, Chang, Majeti and Buenrostro labs for useful discussions. We acknowledge the C. Bustamante lab for help with sequencing.

## Author Contributions

J.D.B., M.R.C., H.Y.C, and W.J.G. conceived the project. M.R.C. and R.M. performed cell sorting. J.D.B. performed ATAC-seq and scATAC-seq data analysis, and oversaw scATAC-seq library generation and protocol optimization performed by B.W. B.W. generated the scATAC, bulk ATAC-seq data and RNA-seq data, M.R.C. performed the RNA-seq data analysis. A.N.S developed the TF motif analysis tools and C.A.L. developed the associated web resources. J.D.B. and W.J.G. wrote the manuscript with input from all authors.

## Supplementary Materials

Supplementary Table 1: Immunophenotypes and cell isolation

Supplementary Table 2: Quality metrics for scATAC-seq data

Supplementary Data Table 1: TF z-scores and PCA values for each cell

## Supplementary URLs

UCSC genome browser track hub, bulk data: https://s3.amazonaws.com/JasonBuenrostro/scATAC_heme_label/hub.txt

UCSC Genome Browser Track Hub, clusters: https://s3.amazonaws.com/JasonBuenrostro/scATAC_heme/hub.txt.

Web application for visualizing PCA projection and TF z-scores: http://schemer.buenrostrolab.com/.

## Materials and Methods

### Cell collection and isolation

Human bone marrow was sorted as described previously^17^. In addition to the cell types previously described, we also isolated plasmacytoid dendritic cells (pDCs) and megakaryocytes. Megakaryocytes were isolated using *in vitro* differentiation of bone marrow derived CD34^+^ cells to megakaryocytes. To do this, CD34+ cells were cultured in StemSpan SFEM with Megakaryocyte Expansion Supplement (Stem Cell Technologies) for 14 days, yielding approximately 100-fold expansion in cell number. To isolate pDCs from human bone marrow using FACS we gated for live, lineage negative, CD34+ CD38+ CD10- CD45RA+ CD123+ cells. To isolate UNK cells from human bone marrow by FACS, we gated for live, lineage negative, CD34+ CD38+ CD10- CD45RA+ CD123-. After cell isolation using FACS, 15,000 single-cells were resuspended in 100 μl of BAMBANKER™ serum-free cell freezing medium (Wako Chemicals, 302-14681) and cryopreserved in liquid nitrogen.

### Single-cell ATAC-seq

Single-cells not cryopreserved after FACS (fresh) were assayed as previously described^7^. To assay cells cryopreserved after FACS (frozen), cells were allowed to recover for 10 min at 37°C in IMDM with 10% FBS. After recovery, cells were washed twice in cold 1x PBS and once in with the C_1_ DNA Seq Cell Wash Buffer (Fluidigm). Cells were then resuspended in 6 μl of C_1_ DNA Seq Cell Wash Buffer, and were combined with 4 μl of C_1_ Cell Suspension Reagent, 7 μl of this cell mix was loaded onto the Fluidigm IFC. Single-cells were then assayed using scATAC-seq as previously described^7^.

### Bulk ATAC-seq and RNA-seq

ATAC-seq and RNA-seq libraries were generated as previously described^7,17^ with slight modifications for frozen cells. One vial of 15,000 frozen cells in 100 μl of BAMBANKER™ freezing medium was quickly thawed at 37°C. 70 μl for bulk ATAC-seq and 30 μl for bulk RNA-seq were then added to 500 μl of warm IMDM with 10% FBS. For bulk ATAC-seq, the cells were split into 2 tubes of 5,000 cells used as technical replicates. Cells were washed twice in 1x PBS, all supernatant was carefully removed without disturbing the cell pellet, and cells were resuspended in 40 μl of transposition mix (20 μl of 2x TD buffer, 2 μl of TDE1, 0.2 μl of 2% digitonin, 13.33 μl of 1x PBS, and 4.47 μl of nuclease-free water) (Illumina, FC-121-1030; Promega, G9441), here the transposition reactions were scaled down to compensate for cell loss during washes. Transposition reactions were incubated at 37 °C for 30 min in an Eppendorf ThermoMixer with agitation at 300 rpm. The transposed DNA fragments were purified and amplified as described^17^.

For bulk RNA-seq, cells were split into 2 technical replicates. RNA was isolated using the Qiagen RNeasy Plus Micro kit, and RNA was eluted in 10 μl of RNase-free water. 5 μl of total RNA was used as input for NuGen Ovation V2 cDNA synthesis kit. The yield of purified SPIA-amplified cDNA was measured using Qubit dsDNA HS Assay kit. 50 ng of SPIA cDNA was fragmented using Nextera DNA library preparation kit (Illumina, FC-121-1030). Fragmented SPIA cDNA was then purified using Qiagen MinElute Reaction Cleanup Kit, and purified DNA was eluted in 10 μl of elution buffer (10mM Tris-HCl, pH 8). Purified SPIA cDNA fragments was amplified and purified as previously described for ATAC-seq^17^. ATAC-seq and RNA-seq libraries were quantified using qPCR, amplified libraries were sequenced using paired-end, dual-index sequencing on a NextSeq 500 instrument with 76 bp read lengths.

### Data pre-processing and TF scores

Single-cell and bulk ATAC-seq alignment, quality filtering and peak calling was performed as previously described^17^, with one exception. For single-cell profiles, any read-pair that occurred in two cells from the same experiment (a single 96-cell IFC) was removed from further analysis. Using the previously described approach^17^, we defined a peak list using all bulk hematopoietic data analyzed here, resulting in 491,437 500bp non-overlapping peaks which we use for the reminder of this study. To count the number of fragments per peak in a sample or cell, and also for computing TF z-scores per cell, we used the default settings in the chromVAR package (Schep *et al.*). Bulk counts were normalized as described previously^17^ using quantile normalization and the CQN package.

### PCA projection and clustering cells

To calculate PCs on the bulk data sets, used or projecting single-cell profiles, we first removed peaks in annotated promoters or aligning to chrX, peaks associated with chrY and unmapped contigs were filtered out in the preprocessing steps. 455,057 peaks remained after filtering and were used for the PCA projection analysis. To normalize the bulk count matrix by library size, we identified 19,287 low variance promoters across all bulk samples and normalized each sample by the mean fragment counts within the low variance promoters. We subsequently took the mean counts of all normalized bulk sample replicates (HSC, MPP, LMPP, CMP, GMP, MEP, Mono, CD4, CD8, NK, NKT, B, CLP, Ery, UNK, pDC and Megakaryocyte) and performed PCA-SVD, resulting in 15 principal components.

To score single-cells by the activity for each component, we first centered the counts for each cell by dividing each peak by the mean fragment counts in peaks for a given cell. We then used the weighted coefficients for each peak and PC (determined using PCA-SVD of the bulk data above) to take the product of the weighted PC coefficients by the centered count values for each cell, taking the sum of this value resulted in a matrix of cells by PCs. Last, we calculate a cell-cell similarity matrix using Pearson correlation and perform PCA on the similarity matrix of correlation values. To validate the computational approach we repeated this procedure for the bulk data, down sampled bulk data, mean single-cell profiles and synthetic mixtures. Notably, we found a patient-specific batch effect in the HSC single-cell profiles in the PCA projected subspace. The batch signal was strongly correlated with Jun/Fos TF z-scores. We therefore normalized each HSC batch to the mean values of all HSC profiles, and in addition, blacklisted all motifs correlated with Jun/Fos activity. We use these corrected PC values and filtered TF motifs for all subsequent visualizations of these data.

To determine significant cell-cell variability in the PC projected sub-space, we either down sampled or permuted peaks by their GC content and mean accessibility. To permute peaks that match GC% and mean accessibility, we take the sum accessibility of all cells of a given immunophenotypically defined cell type (e.g. HSCs) and use the ChromVAR function “get_background_peaks” with the default settings. Notably, ChromVAR samples background peaks with replacement and may select the observed peak as a background peak, and therefore provides a conservative estimation of excess variability.

### Hierarchical clustering, K-medoids and computing density

All hierarchical clustering performed used Pearson correlation as the distance function. All k-medoids clustering shown in this work is performed using 10 replicates with the distance function Pearson correlation. The gap statistic is used to determine the appropriate number of clusters, here the optimal cluster number is determined wherein the minimum K satisfies the follow criteria: the gap value for a given K is greater than the gap value of K+1 minus the standard error of the clustering solution for K+1. The first 5 PCs were chosen for clustering cells by their projected PC values. All metrics of data density shown in this work are weighted by the expected *in vivo* frequency of each cell type, as measured from flow cytometry data. 3D data density is calculated as the mean number of cells within a radius of 1, weighted by the *in vivo* frequency. 2D data density is calculated using KDE weighted by the *in vivo* frequency.

### Lineage bias analysis

To compute TF variability within epigenome and immunophenotype pure (EIPP) cells, we first determined the dominant cell type for each of the 14 k-medoids determined clusters. We then collected 14 cluster-pure and immunophenotype-pure profiles and proceeded to compute variability and TF z-scores using ChromVAR (Schep *et al.*). To determine the magnitude and direction of the TF lineage bias, we first partitioned TF z-scores as greater than zero (high) or less than zero (low). We then computed the mean of the first 5 PCs from the PCA projection for the cells assigned to the high or low TF z-score, distance of the high and low centroids was calculated using Euclidean distance. We determined significance using "direction z-scores", whereby we repeated the analysis described above for PCs calculated using 50 background peak permutations matching GC and mean accessibility, peaks were determined using ChromVAR (Schep *et al.*). Direction z-scores were computed comparing the observed to the 50 permuted distances.

### Identifying significantly differential CMP peaks

To determine differential CMP peaks, aggregate CMP profiles for each k-medoids cluster were collected. Pairwise binomial tests were performed for each aggregate profile (4 CMP clusters) for each peak. Peaks with a p-value of <10^-5^ in one or more comparisons was used for further analysis (n = 1,801 peaks). For clustering and visualization, counts were normalized using column z-scores and clustered using k-medoids, the gap statistic (as described above) was used to determine a K of 6.

### Ordering cells in myeloid pseudo-time and determining smoothed scores

To order cells by myeloid development, only cells within clusters K1, K2, K9, K10, K11 were considered. To determine a continuous trajectory we first fit a line through each cluster centroid using a linear interpolation across the 5 PCs. To assign each cell to the linear trajectory, we determined the closest point of each cell to the interpolated fit line. To determine the regulatory activity dynamics across this inferred trajectory, we smoothed the TF z-scores along myeloid progression with a lowess function with a span of 200, implemented within the MATLAB function smooth. Continuous profiles were normalized by their min/max value for plotting and downstream clustering. To determine 95% confidence intervals we sampled cells (n=100 permutations) with replacement and repeated the smoothing for each permutation. Accessible peaks were similarly smoothed using a span of 500.

### Filtering for variable TFs, regulatory elements and determining motif enrichment

To determine TFs that were significantly variable within the smoothed myeloid trajectories, we compared the standard deviation of the observed smoothed scores by a set of similarly smoothed, permuted, TF scores by randomly permuting the myeloid cell order. Selecting TFs with a standard deviation greater than 0.5 provided 205 motifs with an FDR < 1%. To determine highly variable regulatory elements we first filtered for regulatory elements with greater than a mean accessibility of 0.01 fragments per cell and subsequently filtered for regulatory elements with a coefficient of variation of greater than 0.5 (n = 3,403).

To determine motif enrichment across dynamic accessible peaks, we first collected log-PWM scores per peak using ChromVAR(Schep *et al.*) for motifs that were selected in the TF analysis described above. Peak-to-peak distances were computed using Euclidean distance of the PC scores determined by PCA of the min/max normalized accessibility dynamics across the myeloid trajectory. For each peak, the mean log-PWM score was computed for the nearest 300 peaks.

## References

1. Waddington, C. The Strategy of the Genes: a discussion of some aspects of theoretical biology. The American Biology Teacher 22, 47–47 (1960).

2. Goldberg, A. D., Allis, C. D. & Bernstein, E. Epigenetics: A Landscape Takes Shape. Cell 128, 635–638 (2007).

3. Calo, E. & Wysocka, J. Modification of Enhancer Chromatin: What, How, and Why? Molecular Cell 49, 825–837 (2013).

4. Long, H. K., Prescott, S. L. & Wysocka, J. Ever-Changing Landscapes: Transcriptional Enhancers in Development and Evolution. Cell 167, 1170–1187 (2016).

5. Takahashi, K. & Yamanaka, S. Induction of pluripotent stem cells from mouse embryonic and adult fibroblast cultures by defined factors. Cell 126, 663–676 (2006).

6. Graf, T. & Enver, T. Forcing cells to change lineages. Nature 462, 587–594 (2009).

7. Buenrostro, J. D. et al. Single-cell chromatin accessibility reveals principles of regulatory variation. Nature 523, 486–490 (2015).

8. Cusanovich, D. A. et al. Epigenetics. Multiplex single-cell profiling of chromatin accessibility by combinatorial cellular indexing. - PubMed - NCBI. Science 348, 910–914 (2015).

9. Jin, W. et al. Genome-wide detection of DNase I hypersensitive sites in single cells and FFPE tissue samples. (2015). doi:10.1038/nature15740

10. Kind, J. et al. Genome-wide Maps of Nuclear Lamina Interactions in Single Human Cells. Cell 163, 134–147 (2015).

11. Smallwood, S. A. et al. Single-cell genome-wide bisulfite sequencing for assessing epigenetic heterogeneity. 11, 817–820 (2014).

12. Rotem, A. et al. Single-cell ChIP-seq reveals cell subpopulations defined by chromatin state. Nat. Biotechnol. 33, 1165–1172 (2015).

13. Orkin, S. H. & Zon, L. I. Hematopoiesis: An Evolving Paradigm for Stem Cell Biology. Cell 132, 631–644 (2008).

14. Becker, A. J., McCulloch, E. A. & Till, J. E. Cytological demonstration of the clonal nature of spleen colonies derived from transplanted mouse marrow cells. Nature 197, 452–454 (1963).

15. Goardon, N. et al. Coexistence of LMPP-like and GMP-like Leukemia Stem Cells in Acute Myeloid Leukemia. Cancer Cell 19, 138–152 (2011).

16. Chen, L. et al. Transcriptional diversity during lineage commitment of human blood progenitors. 345, 1251033–1251033 (2014).

17. Corces, M. R. et al. Lineage-specific and single-cell chromatin accessibility charts human hematopoiesis and leukemia evolution. Nature Genetics 48, 1193–1203 (2016).

18. Novershtern, N. et al. Densely interconnected transcriptional circuits control cell states in human hematopoiesis. Cell 144, 296–309 (2011).

19. Lara-Astiaso, D. et al. Chromatin state dynamics during blood formation. Science 55, 1–10 (2014).

20. Ji, H. et al. Comprehensive methylome map of lineage commitment from haematopoietic progenitors. 467, 338–342 (2010).

21. Manz, M. G., Miyamoto, T., Akashi, K. & Weissman, I. L. Prospective isolation of human clonogenic common myeloid progenitors. Proceedings of the National Academy of Sciences of the United States of America 99, 11872–11877 (2002).

22. Corces-Zimmerman, R. & Buenrostro, J. D. Lineage-specific and single cell chromatin accessibility charts human hematopoiesis and leukemia evolution. 1–58 (2015).

23. Lawrence, H. J. et al. Mice bearing a targeted interruption of the homeobox gene HOXA9 have defects in myeloid, erythroid, and lymphoid hematopoiesis. Blood 89, 1922–1930 (1997).

24. Magnusson, M., Brun, A. C. M., Lawrence, H. J. & Karlsson, S. Hoxa9/hoxb3/hoxb4 compound null mice display severe hematopoietic defects. Experimental Hematology 35, 1421.e1–1421.e9 (2007).

25. Naik, S. H. et al. Diverse and heritable lineage imprinting of early haematopoietic progenitors. Nature 496, 229–232 (2013).

26. Busch, K. et al. Fundamental properties of unperturbed haematopoiesis from stem cells in vivo. Nature 518, 542–546 (2015).

27. Perié, L., Duffy, K. R., Kok, L., de Boer, R. J. & Schumacher, T. N. The Branching Point in Erythro-Myeloid Differentiation. Cell 163, 1655–1662 (2015).

28. Paul, F. et al. Transcriptional Heterogeneity and Lineage Commitment in Myeloid Progenitors. Cell 163, 1663–1677 (2015).

29. Notta, F. et al. Distinct routes of lineage development reshape the human blood hierarchy across ontogeny. Science 1–16 (2015).

30. Fulco, C. P. et al. Systematic mapping of functional enhancer-promoter connections with CRISPR interference. Science 354, aag2445–773 (2016).

31. Buenrostro, J. D., Giresi, P. G., Zaba, L. C., Chang, H. Y. & Greenleaf, W. J. Transposition of native chromatin for fast and sensitive epigenomic profiling of open chromatin, DNA-binding proteins and nucleosome position. Nat. Methods 10, 1213–1218 (2013).

32. Buenrostro, J. D., Wu, B., Chang, H. Y. & Greenleaf, W. J. ATAC-seq: A Method for Assaying Chromatin Accessibility Genome-Wide. Curr Protoc Mol Biol 109, 21.29.1–9 (2015).

33. Consortium, R. E. et al. Integrative analysis of 111 reference human epigenomes. Nature 518, 317–330 (2015).

34. Guo, M. H. et al. Comprehensive population-based genome sequencing provides insight into hematopoietic regulatory mechanisms. Proceedings of the National Academy of Sciences of the United States of America 201619052 (2016). doi:10.1073/pnas.1619052114

35. McKenna, A. et al. Whole-organism lineage tracing by combinatorial and cumulative genome editing. Science 353, aaf7907–aaf7907 (2016).

36. Frieda, K. L. et al. Synthetic recording and in situ readout of lineage information in single cells. Nature (2016). doi:10.1038/nature20777

37. Yu, V. W. C. et al. Epigenetic Memory Underlies Cell-Autonomous Heterogeneous Behavior of Hematopoietic Stem Cells. Cell 167, 1310–1322.e17 (2016).

38. John, S. et al. Chromatin accessibility pre-determines glucocorticoid receptor binding patterns. Nature Genetics 43, 264–268 (2011).

